# A patchwork pathway of apparently recent origin enables degradation of the synthetic buffer compound TRIS in bacteria

**DOI:** 10.1101/2023.08.01.551466

**Authors:** Johannes Holert, Aron Borker, Laura Nübel, Rolf Daniel, Anja Poehlein, Bodo Philipp

**Author notes:** Corresponding authors: Johannes Holert, Bodo Philipp.

## Abstract

The widely used synthetic buffer compound TRIS was long considered to be biologically inert. Herein, we describe the discovery of a complete bacterial degradation pathway for TRIS. By serendipity, a *Pseudomonas* strain was isolated from sewage sludge that was able to grow with TRIS as only carbon and nitrogen source. Genome and transcriptome analyses revealed two adjacent gene clusters embedded in a mobile genetic element on a conjugative plasmid to be involved in TRIS degradation. Conjugational transfer of this plasmid into *P. putida* KT2440 enabled this strain to grow with TRIS, demonstrating that the complete TRIS degradation pathway can be transmitted by horizontal gene transfer. Heterologous gene expression revealed cluster I to encode a TRIS uptake protein, a TRIS alcohol dehydrogenase, and a TRIS aldehyde dehydrogenase, catalyzing the oxidation of TRIS into 2-hydroxymethylserine. Gene cluster II encodes a methylserine hydroxymethyltransferase and a D-serine dehydratase which plausibly catalyze the conversion of 2-hydroxymethylserine into pyruvate. Subsequent enrichments from wastewater purification systems led to the isolation of further TRIS-degrading bacteria from the *Pseudomonas* and *Shinella* genera carrying highly similar TRIS degradation gene clusters.

Our data indicate that TRIS degradation evolved recently via gene recruitment and enzyme adaptation from multiple independent metabolic pathways and database searches suggest that the TRIS degradation pathway is now globally distributed. Our study illustrates how engineered environments can enhance the emergence of new microbial metabolic pathways in short evolutionary time scales. This knowledge is valuable for developing new water purification processes in times of increasing water scarcity.

## Introduction

Since the beginning of the industrial revolution hundreds of thousands of new pesticides, herbicides, medical drugs, and other industrial chemicals were synthesized to meet the growing demand of newly emerging agricultural, pharmaceutical, and industrial applications [1]. Many of these anthropogenic chemicals eventually end up in the environment, where their removal is initially restricted due to the absence of efficient biological degradation mechanisms. However, more and more degradation pathways for chemicals that were introduced in the early twentieth century are now being discovered in microbes, suggesting that functional mineralization pathways can evolve within decades after the initial exposure of the microbial community [2]. A prominent example is the degradation pathway for the anthropogenic herbicide atrazine, which has evolved in the genus *Pseudomonas* around fifty years after the large scale introduction of atrazine to the market, enabling these bacteria to exploit this compound as a carbon and nitrogen source for growth [3]. Similarly, catabolic mineralization pathways for other industrially relevant compounds that were introduced in the early twentieth century including herbicides, pesticides, food additives, and medical drugs are now found in different microorganisms isolated from habitats exposed to these chemicals [4–8]. Here, novel pathways can evolve as a result of sufficient positive selective pressure for either the detoxification of such chemicals or their use as a new nutrient source [2]. Typically, novel degradation pathways emerge in a process called patchwork assembly by the recruitment, adaptation, and mutation of one or more genes encoding promiscuous enzymes from other metabolic pathways, either from within the host organism via gene duplication or via horizonal gene transfer (HGT) from the microbial community [1, 2, 9]. If the mutations happen to increase the substrate affinity and catalytic activity towards the respective new substrate and if the new catalytic activity enables the host to transform the new substrate into metabolites of an already existing metabolic pathway, it will gain a selective advantage over competing microbes by being able to use this compound as a new nutrient and energy resource [1, 2]. The genetic information encoding such novel catabolic pathways is often located on mobile genetic elements, such as plasmids or transposons, enabling transfer of the novel pathway to other microbes via HGT [4, 10, 11]. Eventually, positive selection for the new catabolic trait can lead to a loss of mobility and manifestation of the new pathway in the genome of its host.

However, the environmental concentrations of synthetic compounds may be too low to provide enough selection pressure for the evolution of novel catabolic pathways. Especially in wastewater treatment plants where the availability of other, readily degradable organic compounds is high, many low-concentrated synthetic compounds presumably reside too shortly for the evolution of novel degradation pathways and remain in the purified wastewater as so-called micropollutants [12, 13]. Even some synthetic and natural compounds for which efficient biodegradation pathways exist, such as steroid hormones, are sometimes not fully metabolized in wastewater treatment plants due to their low concentration and contribute to the micropollutant pool. To enhance the biodegradation of such micropollutants, bioaugmentation of water treatment facilities is increasingly being considered, which implies the establishment of microorganisms in the water purification processes, which have been isolated for their ability to efficiently degrade the respective organic compounds [14, 15].

The study presented here was originally set up to isolate bacteria capable of degrading steroid hormones in low concentrations from activated sludge. To increase the chances of isolating strains that can later be established for bioaugmentation in real wastewater treatment plant conditions, we used an artificial wastewater medium to mimic typical wastewater nutrient conditions. Surprisingly, our enrichment procedure resulted in the isolation of a bacterial strain that did not grow with the provided steroid hormones, but with the synthetic medium buffer TRIS (2-amino-2-(hydroxymethyl)propane-1,3-diol). Herein, we pursued this finding further and present the identification and characterization of a bacterial degradation pathway for TRIS, which has been considered to be biologically inert so far.

## Results

### Isolation and characterization of the TRIS-degrading strain *Pseudomonas hunanensis* Teo1

The first TRIS-degrading organism was isolated from an enrichment culture that was originally designed to isolate testosterone-degrading bacteria from activated sludge (**Supp. Methods**). We repeatedly observed growth in the enrichment cultures in TRIS-buffered artificial wastewater medium without removal of the provided carbon source testosterone (not shown). Plating of the enrichment cultures onto solid medium containing testosterone resulted in the isolation of a bacterial strain that was not able to grow with testosterone but showed growth in liquid artificial wastewater medium without the addition of any extra carbon or nitrogen source. Subsequent growth experiments in phosphate buffered minimal medium with different concentrations of TRIS confirmed a linear relationship between the provided TRIS concentration and the amount of biomass produced by the isolate (**Fig. 1A**), suggesting that TRIS was the growth substrate in these cultures. Growth experiments coupled with HPLC-MS analysis confirmed that TRIS was completely removed from the culture medium, suggesting its utilization as a carbon and nitrogen source for growth (**Fig. 1B-D**). The isolate had an exponential growth rate of 0.36 +/− 0.09 h^-1^ and a doubling time of 0.44 +/− 0.09 h (n = 6). The only detectable metabolite accumulating in the medium was ammonium, with around 40 % of the nitrogen introduced with the TRIS substrate being released as ammonium into the medium (**Fig. 1C-D**). The strain did not grow with the structurally similar amino alcohols serinol, 2-amino-2-methyl-1-propanol, and 2-amino-2-methyl-1,3-propanediol (**Supp. Fig. S1**), while it did grow with the quaternary ammonium compounds choline and betaine, and the secondary amine sarcosine, and with the amino acids D- and L-serine, but not with glycine. Based on its whole genome sequence (**Supp. Methods**), the isolate was taxonomically classified as a representative of the species *Pseudomonas hunanensis* and was named strain Teo1 (**Supp. Tab. ST1**). The genome of strain Teo1 consists of a circular chromosome (6.13 Mbp) and two large, circular plasmids, p1_Teo1 (156.3 kbp) and p2_Teo1 (65.0 kbp).

**Fig. 1:**
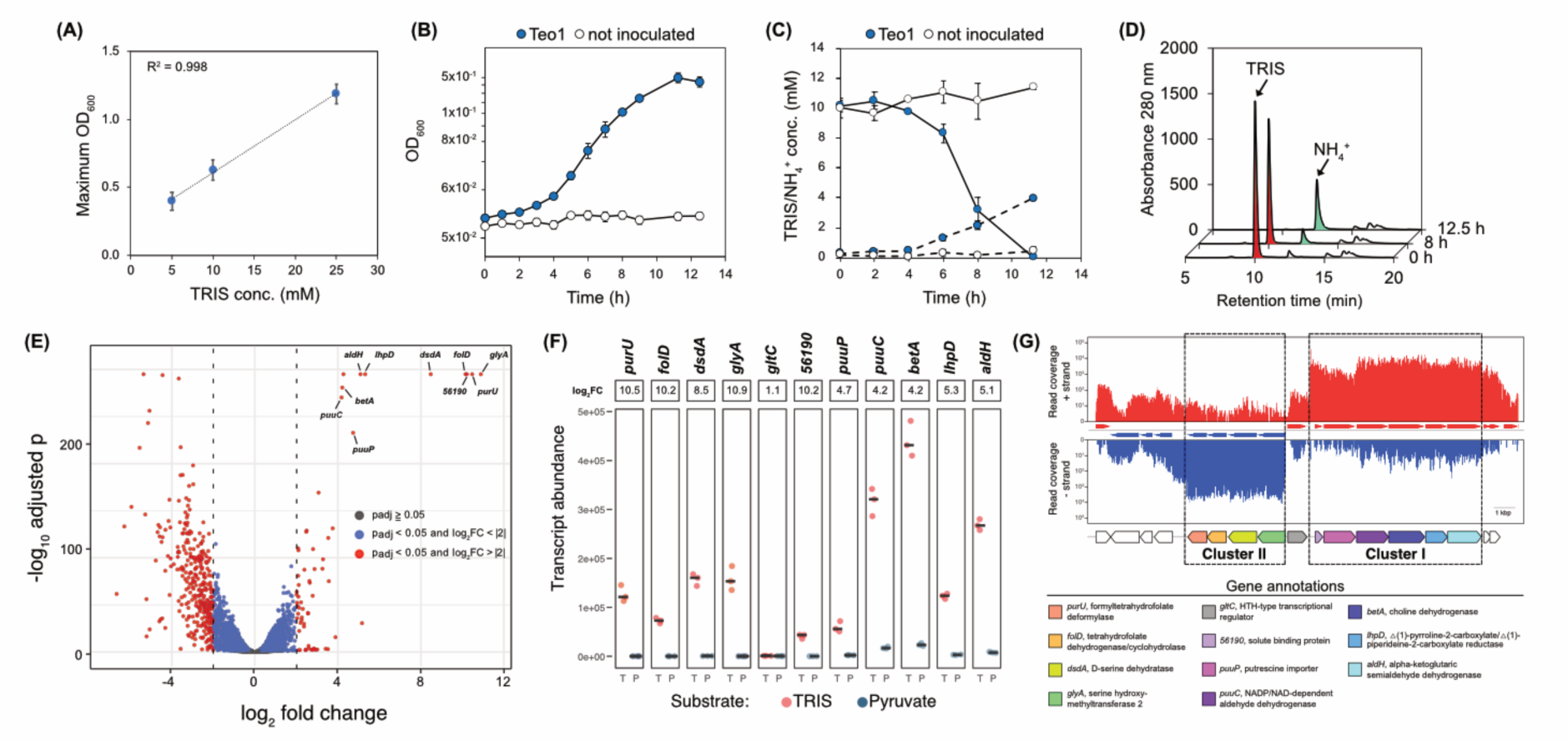
Characteristics of TRIS metabolism in *Pseudomonas hunanensis* Teo1 and identification of potential TRIS degradation genes. (**A**) Maximal biomass production of strain Teo1 in relation to TRIS concentration. Error bars indicate standard deviation (n = 3). (**B**) Growth of strain Teo1 with 10 mM TRIS as only carbon and nitrogen source (n = 6). (**C**) Removal of TRIS (straight lines) and formation of ammonium (dashed lines) in Teo1 growth cultures. (**D**) Representative HPLC chromatograms of a strain Teo1 culture growing with 10 mM TRIS. TRIS is depleted from the culture while ammonium is released. (**E**) Volcano plot of transcriptome data showing significantly up-regulated genes (red) under TRIS-grown (positive fold changes) and pyruvate-grown conditions (negative fold changes) in Teo1. Genes predicted to be involved in TRIS degradation are labelled. (**F**) Transcript abundances and log2(fold changes) of the predicted TRIS degradation genes in Teo1 in TRIS- and pyruvate-grown cells (three biological replicates of each growth substrate, horizontal bar indicates median, corrected p values of all selected genes were < 0.05). (**G**) mRNA reads (red: positive strand, blue: negative strand) of a representative TRIS-grown culture mapped onto the DNA section containing the predicted TRIS degradation gene clusters in strain Teo1.

### Identification of TRIS degradation genes

TRIS growth experiments and time-resolved analysis of TRIS degradation in cell suspensions of strain Teo1 showed that Tris degradation was induced in cells pre-grown with TRIS while it was not induced in pyruvate pre-grown cells (**Supp. Fig. S2**), suggesting that TRIS catabolism is regulated at the transcriptional level in strain Teo1. Based on this observation we used transcriptomics of TRIS-grown and pyruvate-grown Teo1 cells to identify potential TRIS degradation genes. A total of 398 genes were differentially regulated in TRIS-compared to pyruvate-grown cells (fold change (FC) > 4, padj < 0.05, **Fig. 1E**, **Dataset SD1**), of which 65 were upregulated in TRIS-grown cells. Ten of the twelve most upregulated genes in TRIS-grown cells were found in two adjacent gene clusters (**Fig. 1E-G**) localized on the p1_Teo1 plasmid. The proteins encoded in cluster I were annotated as a solute binding protein, an amino acid/polyamine transporter, a choline dehydrogenase, a NAD/NADP-dependent aldehyde dehydrogenase, a △(1)-pyrroline-2-carboxylate/△(1)-piperideine-2-carboxylate reductase, and an alpha-ketoglutaric semialdehyde dehydrogenase (**Tab. 1**). The proteins encoded in cluster II were annotated as a serine hydroxymethyltransferase, a D-serine dehydratase, and two proteins presumably involved in the transformation of methylene tetrahydrofolate into tetrahydrofolate and formate. The clusters are separated by a gene that encodes a helix-turn-helix (HTH)-type transcriptional regulator. Based on the similar upregulation of their genes, both gene clusters appear to be expressed as operons (**Fig. 1G**). Due to their strong upregulation and their colocalization in putative operons we proposed that these genes or a subset of them are involved in TRIS degradation in strain Teo1.

**Tab. 1:**
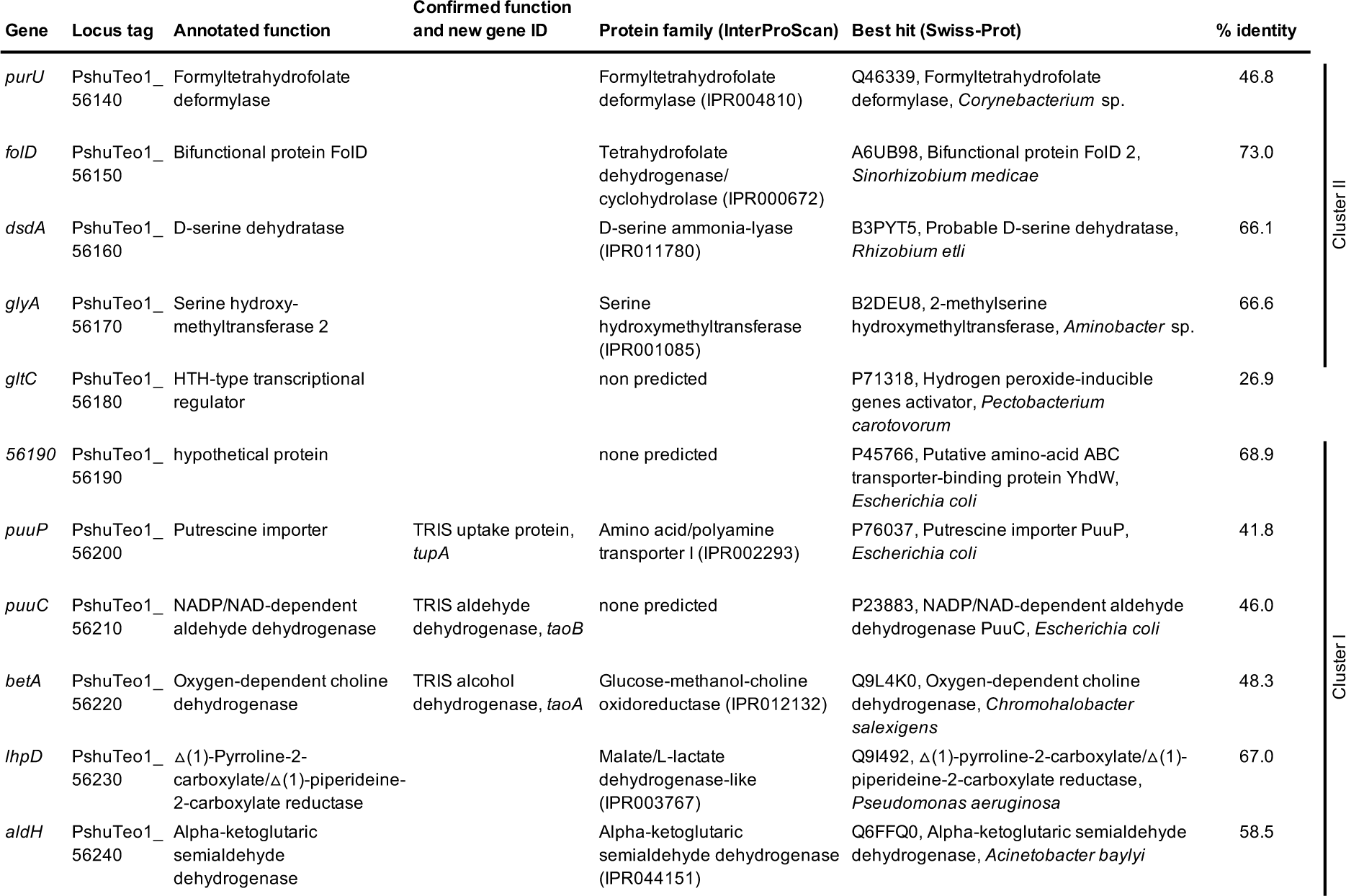
Genes potentially involved in TRIS degradation in *Pseudomonas hunanensis* Teo1.

### Characterization of the initial TRIS degradation reactions

Based on the predicted protein functions encoded in the upregulated gene clusters, we hypothesized that TRIS is initially oxidized by the proteins encoded in cluster I. To test this, we cloned combinations of the genes encoding the putrescine importer-like protein, the choline dehydrogenase-like protein and the aldehyde dehydrogenase into the expression plasmid pUCp18 into *E. coli* (**Suppl. Methods**) and tested high density cell suspensions of the resulting strains for TRIS transformation activity. While the *E. coli* strains that carry only the putrescine importer gene or the choline and aldehyde dehydrogenase genes did not transform TRIS, the strain with all three genes transformed TRIS almost completely into a new compound within 24 h (**Fig. 2A**). This compound was identified as 2-hydroxymethylserine (**Fig. 2B**) based on its molecular weight, which was 14 Da heavier than that of TRIS, suggesting the oxidation of one hydroxymethyl group into a carboxylic group. To test whether this presumable TRIS degradation intermediate can be further utilized by strain Teo1, sterilized culture supernatants containing 2-hydroxymethylserine were used as growth medium for Teo1, confirming that strain Teo1 can grow with and fully degrade 2-hydroxymethylserine (**Fig. 2C**). These results supported our hypothesis that TRIS is transported into the cells by the putrescine importer-like protein and subsequently oxidized by the choline dehydrogenase-like protein and the aldehyde dehydrogenase. Thus, we propose that these genes encode a TRIS uptake protein (TupA), and two TRIS-oxidizing proteins, namely a TRIS alcohol dehydrogenase (TaoA) and a TRIS aldehyde dehydrogenase (TaoB).

**Fig. 2:**
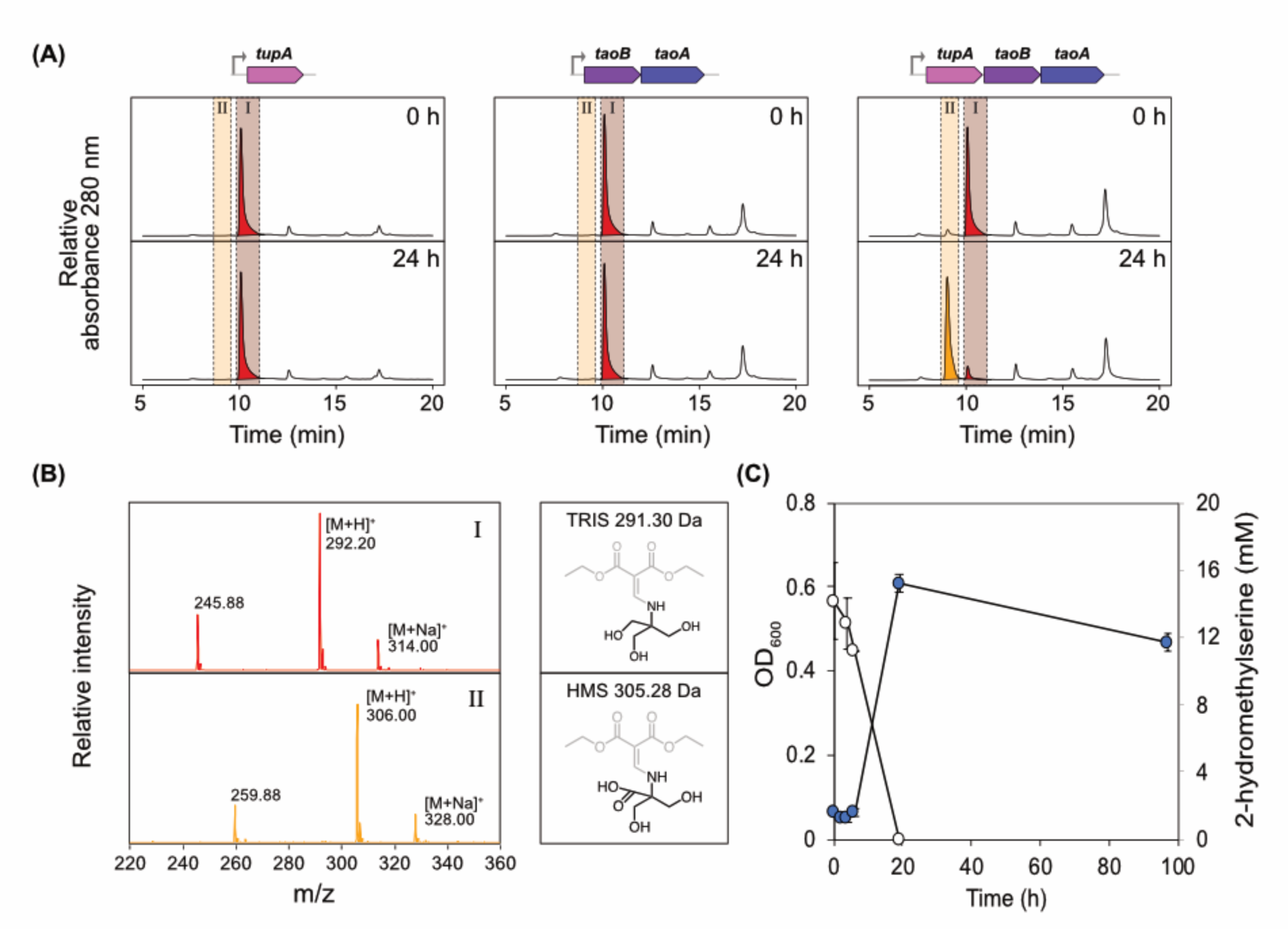
TRIS transformation activity of *E. coli* strains carrying different combinations of the putrescine importer-like protein-encoding gene (*puuP,* now *tupA*), the aldehyde dehydrogenase-encoding gene (*puuC,* now *taoB*), and the choline dehydrogenase-like protein-encoding gene (*betA,* now *taoA*) on the expression plasmid pUCp18. (**A**) Representative HPLC-UV/Vis chromatograms of TRIS-containing cell suspension supernatants of *E. coli* strains that carried either the *tupA* gene, the *taoB* and *taoA* genes, or all three genes. TRIS (peak I, red) was transformed into a new compound (peak II, orange) in the *E. coli* strain that carries all three genes only. The cloned genes were under the activity of the *lac* promotor (grey arrow). (**B**) Mass spectra of TRIS (peak I) and the new product (peak II). The new product was identified as 2-hydroxymethylserine based on the molecular mass of its diethyl ethoxymethylenemalonate (DEEMM, grey molecule moiety) derivative of 305 Da, which is 14 Da heavier than the molecular mass of 291 Da of the DEEMM derivative of TRIS. (**C**) Growth of strain Teo1 (blue circles) in sterilized culture supernatants containing 2-hydroxymethylserine produced from TRIS and removal of 2-hydroxymethylserine (white circles). Average and standard deviation of three independent biological replicates are shown.

### Isolation of further TRIS-degrading bacteria and genomic analysis of TRIS degradation

Subsequent targeted enrichment of TRIS-degrading bacteria resulted in the isolation of seven *Pseudomonas* strains (Teo2, Teo3, Teo4, Teo6, Teo8, Teo10, and Teo11) and one *Shinella zoogloeoides* strain (Teo12) (**Supp. Tab. ST1**). All isolates were all able to grow with TRIS as only carbon and nitrogen source and completely removed TRIS from the culture medium (**Supp. Fig. S3**). The *Pseudomonas* strains Teo2, Teo4, Teo6, Teo8, and Teo11 and the *Shinella* isolate strain Teo12 were isolated from different municipal wastewater treatment plant samples, *Pseudomonas sichuanensis* Teo3 from an activated charcoal basin of a water purification plant, and *Pseudomonas* sp. strain Teo10 from a freshwater lake sample. Enrichment cultures from two additional wastewater treatment plants and from five freshwater habitats did not develop growth (**Supp. Tab. ST1)**.

We sequenced the genomes of five *Pseudomonas* isolates (**Supp. Methods**) and found homologous TRIS degradation gene clusters to the ones identified in strain Teo1 in all of these strains (**Fig. 3, Suppl. Fig. S4**). In *Pseudomonas* strains Teo2, Teo4, Teo6, and Teo8 the clusters are also localized on plasmids (p1_Teo2, p1_Teo4, p1_Teo6, and p2_Teo8) and the plasmids have high similarities to p1_Teo1 (**Fig. 3A**). In *Pseudomonas* strain Teo3, the clusters are encoded in the chromosome (**Fig. 3B**). While all gene clusters II are highly similar, only the *tupA*, *taoA*, and *taoB* genes are conserved in all cluster I sequences (**Fig. 3, Suppl. Fig. S4**), suggesting that the △(1)-pyrroline-2-carboxylate/△(1)-piperideine-2-carboxylate reductase-like protein and the alpha-ketoglutaric semialdehyde dehydrogenase-like protein are not required for TRIS degradation. In all these strains, the TRIS degradation gene clusters are flanked by sequences similar to known bacterial insertion elements (**Fig. 3B**), suggesting that the TRIS degradation gene clusters are located within a mobile genetic element [11]. A search in the NCBI nucleotide database identified one *Priestia* genome (accession CP065422.1) and three *Pseudomonas* genomes (CP069081.1, CP097105.1, CP114115.1) with homologous gene clusters **(Fig. 3A, Suppl. Fig. S4)**. In *Priestia*, and two of the *Pseudomonas* strains the clusters are also plasmid-borne, while they are encoded chromosomally in *Pseudomonas* sp. strain GXZC (**Fig. 3B**).

**Fig: 3:**
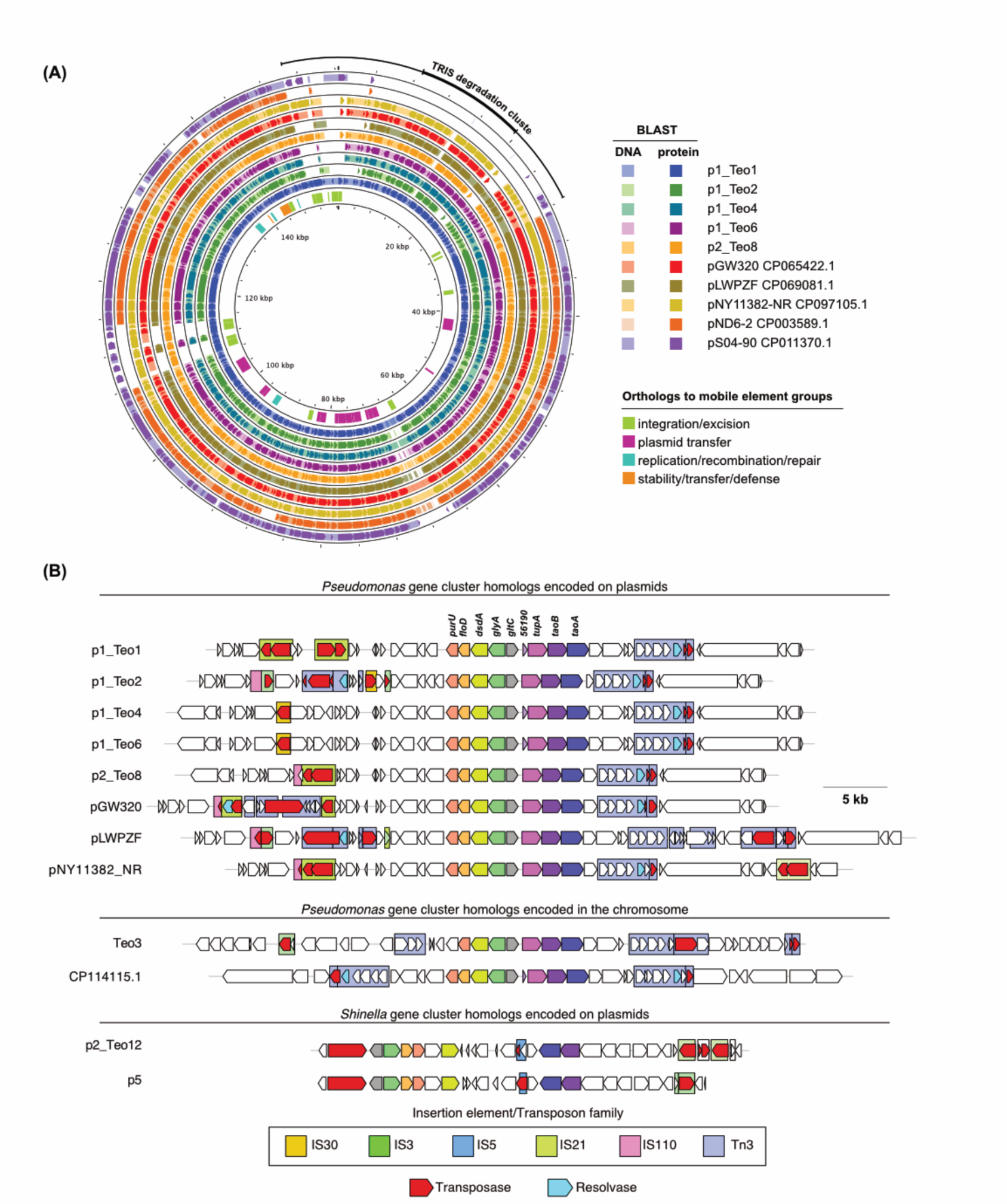
TRIS degradation gene clusters in the genomes of the sequenced TRIS-degrading isolates from this study and of homologous gene clusters found in the NCBI nr nucleotide database. (**A**) Most TRIS degradation gene clusters are located on large circular plasmids with high sequence similarities to the *Pseudomonas* plasmids pND6-2 and pS04-90, which lack the TRIS degradation clusters. The outer black line indicates the location of the putative mobile genetic element including the TRIS degradation genes. Like pND6-2 and pS04-90, the TRIS degradation-encoding plasmids encode orthologs to mobile elements groups (inner circle) including a type IVB secretion system and a type IV pilus for plasmid transfer as well as a type II toxin-antitoxin system. (**B**) All TRIS degradation gene clusters, including plasmid- and chromosome-encoded clusters, are flanked upstream and downstream by transposon- or insertion element-like structures. The plasmid p2_Teo12 from *Shinella* sp. strain Teo12 and the homologous plasmid p5 in a *Rhizobium* strain have no similarity to the other TRIS degradation-encoding plasmids, but also carry two gene clusters which encode homologs to the other TRIS degradation proteins.

We also sequenced the genome of the *Shinella* isolate (**Supp. Methods**), which consists of one circular chromosomes (3.54 Mbp) and three circular plasmids, p1_Teo12 (1.89 Mbp), p2_Teo12 (221.54 kbp) and p3_Teo12 (140.01 kbp), which have no apparent similarity to the TRIS degradation-encoding *Pseudomonas* plasmids. However, a DNA region on p2_Teo12 encodes homologs of all TRIS degradation genes in two adjacent gene clusters, except for the genes encoding the solute binding protein, the uptake protein TupA, the △(1)-pyrroline-2-carboxylate/△(1)-piperideine-2-carboxylate reductase-like protein, and the alpha-ketoglutaric semialdehyde dehydrogenase-like protein (**Fig. 3B, Suppl. Fig. S4**). In these clusters, the *taoA* and *taoB* gene homologs are encoded in the same orientation as in cluster I of the *Pseudomonas* TRIS degradation gene clusters, while the genes are differently oriented in cluster II. A homologous regulator gene to the one found in the *Pseudomonas* gene clusters is located upstream of this cluster. A search in the NCBI nucleotide database identified a closely related *Rhizobium oryzihabitans* strain M15 (CP048637.1), that carries a homologous DNA segment on a plasmid (p5, **Fig. 3B**). The predicted TRIS degradation gene clusters in *S. zoogloeoides Teo12* and strain M15 are separated by a truncated transposase gene and flanked by additional transposase genes and insertion elements.

A phylogenetic analysis of the TaoA proteins with closely related proteins of the glucose-methanol-choline oxidoreductase family (GMC, InterPro IPR012132) revealed that the TaoA proteins form a unique cluster that diverges from a clade containing choline dehydrogenase protein sequences (**Supp. Fig. S5A**). In contrast, other choline dehydrogenase proteins of the TRIS-degrading isolates encoded outside of the TRIS degradation gene clusters cluster within the choline dehydrogenase clade, except for two choline dehydrogenase proteins from *S. zoogloeoides Teo12*, which cluster at the base of the TaoA cluster. A phylogenetic analysis of the TaoB proteins with closely related proteins containing an aldehyde dehydrogenase domain (InterPro IPR015590) showed that the TaoB proteins also from a distinct cluster within the dataset with closest relations to a reviewed aldehyde dehydrogenase (PuuC, Accession P23882) that oxidizes aldehydes during putrescine catabolism in *E. coli* (**Supp. Fig. S5B**). Other PuuC-like proteins encoded outside of the TRIS degradation gene clusters of the TRIS-degrading isolates cluster in close proximity to the TaoB cluster.

### Conjugational transfer of TRIS degradation plasmids

The TRIS degradation-encoding *Pseudomonas* plasmids have high similarities to the previously described large plasmids pND6-2 from *Pseudomonas putida* ND6 and pS04-90 from *Pseudomonas aeruginosa* S04-90 [16, 17]. Both these plasmids lack the putative mobile genetic element that encodes TRIS degradation (**Fig. 3A, Suppl. Fig. S4**). However, pS04-90 carries a carbapenem resistance-encoding class 1 integron at almost the same site in the pND6-2 backbone as the TRIS degradation-encoding elements in the *Pseudomonas* plasmids. Both pND6-2 and pS04-90 are conjugative plasmids that can be transferred from their hosts into other *Pseudomonas* strains. Like pND6-2 and pS04-90, p1_Teo1 and its homologs carry genes encoding type IV pili and a type IVB secretion system (**Fig. 3A, Suppl. Fig. S4**), suggesting that these plasmids are also conjugative plasmids. To test this, we set up conjugation experiments with strain Teo1 and strain Teo8 as plasmid donors and *Pseudomonas putida* KT2440 as a non-TRIS-degrading recipient. To be able to differentiate between donor and recipient cells, we labelled strain KT2440 with a chromosomally integrated gentamycin resistance associated with a yellow fluorescent protein (YFP) gene. After the conjugation of Teo1 and Teo8 with strain KT2440::*eyfp-gm* growth of fluorescing colonies was observed on gentamycin-containing minimal medium plates with TRIS as only carbon and nitrogen source (**Fig. 4A**), suggesting that conjugational transfer of the p1_Teo1 and p2_Teo8 plasmids also transferred the ability to use TRIS as growth substrate to KT2440::*eyfp-gm*. The resulting transconjugants were also able to grow with TRIS in liquid culture (**Fig. 4D**), confirming that both gene clusters together encode the complete degradation pathway sufficient to use TRIS as sole carbon and nitrogen source.

**Fig. 4:**
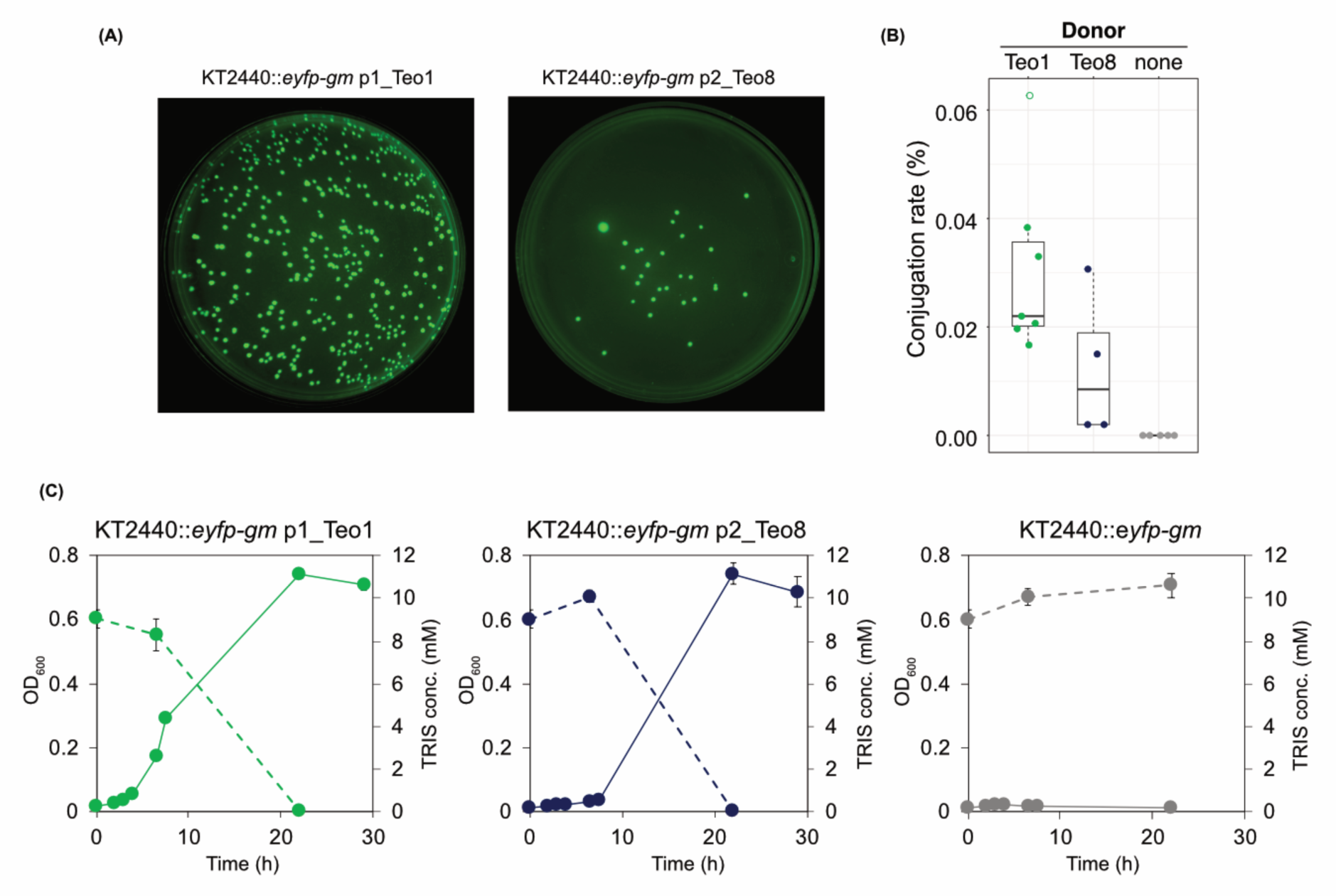
Conjugative transfer of the TRIS degradation pathway-encoding plasmids p1_Teo1 and p2_Teo8. (**A**) Growth of YFP-fluorescing colonies on TRIS-containing agar plates after conjugation of *P. putida* KT2440::*eyfp-gm* with Teo1 (left panel) and Teo8 (right panel) as donor strains. (**B**) Conjugation rate of KT2440::*eyfp-gm* with p1_Teo1, p2_Teo8, and of a control without donor cells. Black lines indicate median rates of three independent conjugation experiments, boxes indicate the interquartile range, and dashed lines the maximum and minimum range. Transconjugants of up to seven replicates from three independent experiments were counted. Datapoints are coloured based on the donor strain. Outliers are not filled. (**C**) Degradation of TRIS (dashed lines) and growth (solid lines) of *P. putida* KT2440::*eyfp-gm* p1_Teo1 and p2_Teo8 transconjugants in liquid medium. The recipient strain *P. putida* KT2440::*eyfp-gm* without the conjugated plasmids was neither able to grow with, nor to degrade TRIS.

## Discussion

In this study we identify and characterize a degradation pathway for the synthetic buffer compound TRIS in several *Pseudomonas* strains and a *Shinella* strain, which were all isolated from wastewater purification systems with TRIS as only carbon source. Our results suggest that TRIS degradation proceeds via a pathway that originated recently via a combination of genes from different, formerly independent metabolic pathways and their assembly into two co-regulated gene clusters within a mobile genetic element. Although not all individual functions for the proteins encoded in clusters I and II were biochemically confirmed, conjugational transfer of the plasmid that contains both clusters into *P. putida* KT2440 co-transferred the ability to degrade TRIS, confirming that they together encode the complete degradation pathway sufficient to use TRIS as sole carbon and nitrogen source.

TRIS has long been considered biologically inert, because the bonding of its amino group to a tertiary carbon atom prevents its metabolism via transamination or oxidative deamination, which are the most common reactions in the degradation of amines. However, bacterial degradation of α-amino acids with a tertiary α-carbon atom is known for a long time and has been demonstrated for 2-hydroxymethylserine via a tetrahydrofolate-dependent methylserine hydroxymethyltransferase (mSHMT) with D-serine and formaldehyde as products [18]. All of our TRIS-degrading isolates encode a protein in cluster II with highest similarities to confirmed D-amino acid-producing mSHMTs [19] as well as a protein for the degradation of D-serine to pyruvate, suggesting that these proteins together can catalyze the degradation of 2-hydroxymethylserine, the product of TRIS oxidation by the TaoAB proteins. Thus, we propose a pathway for the catabolism of TRIS, in which TRIS is translocated into the cytoplasm by the TRIS uptake protein TupA and subsequently oxidized to 2-hydroxymethylserine by the TRIS alcohol dehydrogenase TaoA and the TRIS aldehyde dehydrogenase TaoB via a suspected aldehyde intermediate (**Fig. 5**). 2-Hydroxymethylserine is further transformed into D-serine by the mSHMT protein, which is then transformed into pyruvate by the D-serine dehydratase. The latter can be channelled into the tricarboxylic acid cycle via oxidative decarboxylation to acetyl-CoA. The formaldehyde-accepting tetrahydrofolate (THF) cofactor of the mSHMT protein can then be recycled by the bifunctional methylenetetrahydrofolate dehydrogenase/ methenyltetrahydrofolate cyclohydrolase FolD and the formyltetrahydrofolate deformylase PurU encoded in cluster II [20, 21].

**Fig. 5:**
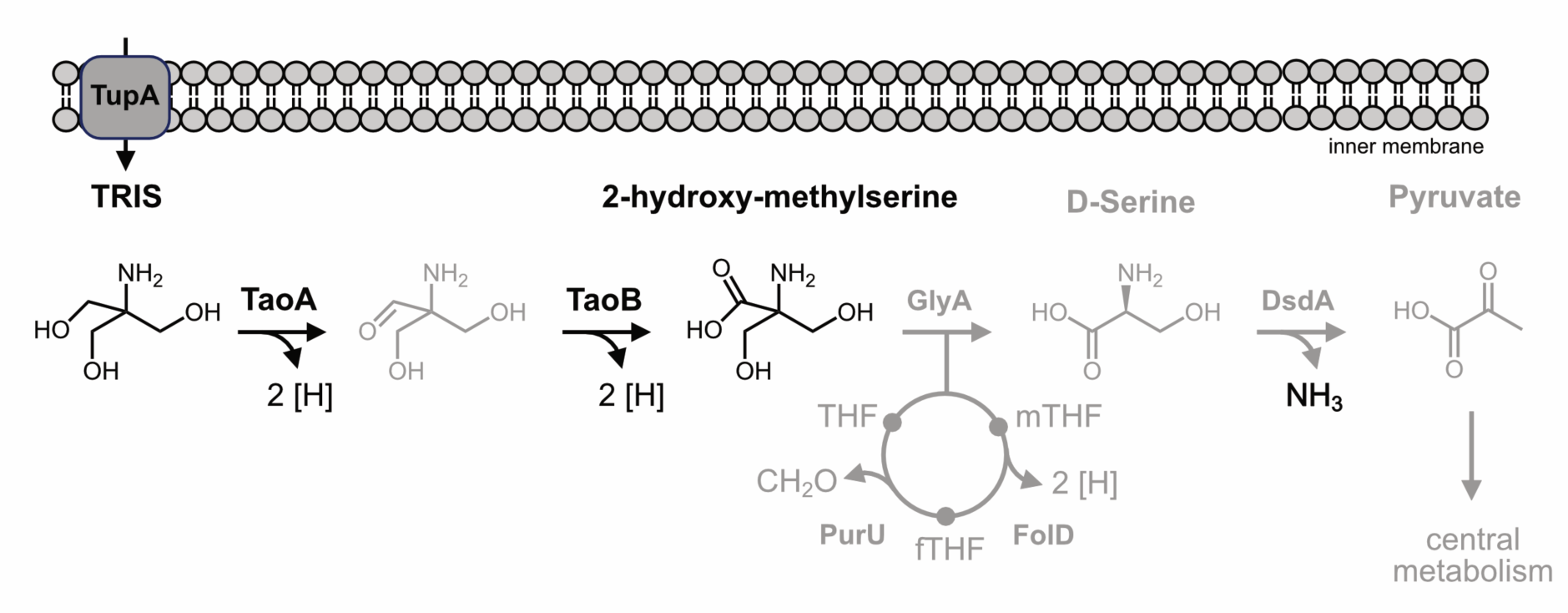
Proposed TRIS degradation pathway. Confirmed reactions and intermediates are in black, proposed reactions and intermediates are in grey, THF = tetrahydrofolate, mTHF = methenyltetrahydrofolate, fTHF = formyltetrahydrofolate.

The genetic context of the *tupA*, *taoA* and *taoB* genes is not found outside of cluster I, suggesting that these genes were assembled from multiple origins. The TaoA proteins have highest similarities to choline dehydrogenases (CDH), which catalyse the oxidation of choline into betaine aldehyde [22] and belong to the glucose-methanol-choline oxidoreductase family (GMC). Within a phylogenetic GMC protein tree, the TaoA proteins form a distinct clade that diverges from the CDH sequences with strong bootstrap support, suggesting that the TRIS alcohol dehydrogenase functionality evolved from enzymes with choline dehydrogenase activity. Strikingly, two CDH proteins encoded outside of gene cluster I in *S. zoogloeoides* Teo12 cluster basal to the TaoA protein clade, suggesting that they could be ancestral to TaoA. It is tempting to speculate that the TaoA proteins have originated in *Shinella* or a similar organism through the duplication and mutation of CDH-encoding genes. TupA and TaoB have highest similarities to proteins involved in the degradation of polyamines like putrescine [23, 24], suggesting that these proteins were recruited from bacterial polyamine degradation pathways [25]. The fact that Teo1 was not able to use other α-substituted amino alcohols such as 2-amino-2-methylpropanol and 2-amino-2-methylproandiol as growth substrates suggests that either the substrate uptake by TupA or the oxidation reactions catalysed by TaoAB evolved to be highly specific for TRIS.

Expression of *tupA* was strictly required for TRIS transformation in *E. coli* cells, suggesting that TRIS translocation is required for TRIS uptake in Gram-negative cells. With a pKa of 8.07 TRIS is mainly positively charged at neutral and acidic pH, preventing its diffusion across the cytoplasmic membrane. The absence of a *tupA* gene homolog in *S. zoogloeoides* Teo12 suggests that this gene was recruited later to the pathway. It remains unknow how TRIS is translocated by Teo12, however, alternative putrescine uptake transporters [23] are also encoded in the genome of Teo12, which might play a role in TRIS uptake in this strain.

The genetic context of the genes encoded in cluster II is also present in other *Pseudomonas* strains that do not encode cluster I, suggesting that this gene cluster may have been assembled prior to its recruitment into the TRIS degradation pathway. While SHMT and mSHMT proteins are often accompanied by THF co-factor recycling proteins [19, 26], their genetic association with D-amino acid degradation genes such as the D-serine dehydratase genes in cluster II in a putative operon was not reported so far. In pathogenic bacteria, D-serine dehydratase genes play important roles in the detoxification and catabolism of D-amino acids [27, 28]. Thus, we conclude that the second part of the TRIS degradation pathway was assembled by a combination of genes involved in the transformation of α-substituted amino acids and in the degradation of D-amino acids. The presence of mostly incomplete insertion elements up- and downstream of the TRIS degradation gene clusters suggests that the whole DNA segment was originally able to be mobilized as a composite transposon, allowing its transposition between plasmids and the chromosome. The finding of an additional transposase gene between the two gene clusters in *Shinella* and the closely related *Rhizobium* strain corroborates our hypothesis that the pathway was assembled from two independent reactions sequences encoded in clusters I and II.

TRIS is used in countless applications in research, industry, medicine, and household settings. Its most prominent applications include its use as a buffer in molecular biology and biochemistry labs, as an excipient in drug formulations, and as an active agent for the treatment of metabolic acidosis and an enhancer for antibiotics and antiseptics for the treatment of Gram-negative infections. Based on its first large scale production in the middle of the twentieth century [29, 30], we hypothesize that the TRIS degradation pathway evolved within the last thirty to fifty years, which is corroborated by the fact that TRIS degradation was not discovered earlier despite its intense use in research laboratories. However, the organization of the TRIS degradation genes in two compact gene clusters and the fact that expression of these presumable operons was regulated in response to TRIS in strain Teo1 are signs for a highly evolved pathway [5], suggesting its turn-around in the environment for a while. The regulator gene between the two clusters is presumably involved in this regulation. Its protein product is similar to the transcription factor GltC that regulates glutamate synthase expression in *Bacillus subtilis* [31]. The first reports of bacteria being able to utilize TRIS as a growth substrate were published thirty years ago from a group in Spain [32], however, no genetic or biochemical analyses followed this observation.

Specific microbial habitats such as sinks and wastewater treatment facilities of large research laboratories, hospitals, or from the TRIS-manufacturing industry could be locations where the TRIS degradation pathway might have originated. First, such habitats encounter significant TRIS concentrations on a regular basis. For example, the antibiotic fosfomycin, which is produced and administered as a TRIS salt, was found at concentrations of 0.13 mM in the influent and 0.015 mM in the effluent of a wastewater treatment plant of a fosfomycin production plant [33], suggesting the presence of similar concentrations of its counterion TRIS. Second, sinks and water purification systems are hotspots for horizontal gene transfer (HGT) [11, 34, 35], which additionally favors the evolution of novel catabolic pathways.

The identification of identical TRIS degradation gene clusters in bacteria that have been isolated in Germany, the Republic of Korea, and in China corroborates our findings, that TRIS degradation-encoding plasmids can be readily transferred among bacteria by conjugation-driven HGT. Our isolation studies suggest that the TRIS degradation pathway is frequently present in the microbiome of municipal wastewater treatment plants but less frequent in freshwater habitats, indicating that environments with high carbon and nitrogen loads better support the maintenance and dissemination of the TRIS degradation pathway than more oligotrophic environments. Although the average TRIS concentration in the sampled wastewater treatment plants is presumably very low, its metabolism as an additional nitrogen and carbon source alongside other recalcitrant molecules might provide sufficient positive selection pressure for bacteria with broad substrate ranges such as *Pseudomonas* and *Shinella* [36–39] to maintain the pathway. Our results corroborate the importance of HGT and mobile genetic elements in the evolution of novel catabolic pathways. With this, they provide additional knowledge about the basic structure of such elements, which can be used for future use in engineering bacteria for bioaugmentation [11].

## Experimental procedures

Additional experimental methods can be found in the supplemental information.

### Targeted enrichment and growth cultures

TRIS-degrading strains were isolated and grown in a phosphate-buffered minimal medium (pH 7.0) containing 0.01 mM CaCl2 x 2 H2O, 1.5 mM MgSO4 x 7 H2O, 35 mM K2HPO4 x 3 H2O, 15 mM NaH2PO4 x H2O, trace elements and 10 mM TRIS. For solid media plates, 1.5 % agar was added prior to autoclaving. Enrichment cultures (5 ml) were inoculated with fresh samples (1 ml) from local wastewater treatment plants (activated sludge or activated carbon), a water purification plant (activated carbon), or different freshwater habitats (**Supp. Tab. ST1**). When enrichment cultures showed growth, 100 µl of the culture supernatant were transferred to 5 ml fresh minimal medium with TRIS, and this was repeated two to three times before cultures were spread onto solid minimal medium plates containing to isolate individual TRIS-degrading colonies. Growth experiments in liquid cultures were carried out at 30°C and 200 rpm in 1 ml or 5 ml medium in 24-well plates or in reaction tubes, respectively. Experimental cultures were inoculated from washed, over-night starter cultures grown with TRIS. Growth was followed by measuring the optical density at 600 nm (OD600). For transcriptome experiments, TRIS was replaced with 20 mM pyruvate and 20 mM NH4Cl were added as nitrogen source. For substrate tests, TRIS was replaced with 10 mM D- or L-serine, 10 mM choline, 10 mM betaine, 10 mM 2-amino-2-methylpropanol, 10 mM 2-amino-2-methylproandiol, 10 mM sarcosine, or 10 mM serinol, and 20 mM NH4Cl were added as nitrogen source. *Pseudomonas putida* KT2440 and its derivatives were grown in the same minimal medium with succinate as carbon source at 30°C. *E. coli* strains were grown in half-concentrated lysogeny broth (LB) at 37°C. If required, kanamycin (20-50 µg/ml), tetracycline (10 µg/ml), or gentamycin (20-40 µg/ml) antibiotics were added to the medium after autoclaving. To test whether 2-hydroxymethylserine can be used as a growth for Teo1, cell suspensions of *tupA*, *taoB* and *taoA* expressing *E. coli* cells that had produced 2-hydroxymethylserine from TRIS were centrifuged and sterile filtered (0.2 µm) and the culture supernatant was inoculated with strain Teo1 cells.

### TRIS quantification

TRIS, its degradation intermediates, and ammonium were identified and quantified by high performance liquid chromatography (HPLC) coupled to a mass spectrometer and a UV/Vis detector [40] after derivatization of their amino group with diethyl ethoxymethylenemalonate (DEEMM, [41]). For this, 100 µl culture supernatants were mixed with 130 µl borate buffer (1 M, pH 9.0), 75 µl methanol, and 3 µl DEEM reagent, followed by 30 min sonication and 60 min incubation at 70°C before injection into the HPLC. A gradient (flow rate 0.3 ml/min) of ammonium-acetate buffer (10 mM, pH 3.0, with 0.1 % (v/v) formic acid, eluent A) and acetonitrile (eluent B) were used for product elution starting with 10 % eluent B for 2 min, increasing to 90 % eluent B within 22 min, remaining at 90 % eluent B for 2 min, and returning to 10 % eluent B within 1 min, followed by an equilibration of 5 min. Products were identified based on their mass spectra after electrospray ionization in positive ion mode. The capillary voltage was 2500 V, the end plate offset 500 V and the capillary temperature 300°C. Nebulizer pressure was 22.5 psi, dry gas flow 12 l/min and dry temperature 200°C. Quantification was performed based on the product peak areas at 280 nm UV absorbance, and concentrations were calculated based on concentration curves of authentic standards treated in the same way. The concentration of 2-hydroxymethylserine was calculated using the same standard curve as for TRIS.

### Transcriptome analysis

*P. hunanensis* Teo1 was grown in minimal medium with TRIS as only carbon and nitrogen source or with pyruvate and ammonium and cultures were harvested during mid-log phase. Harvested cells were re-suspended in 800 μl RLT buffer (RNeasy Mini Kit, Qiagen) with β-mercaptoethanol (10 μl ml^-1^) and cell lysis was performed using a laboratory ball mill. Subsequently, 400 μl RLT buffer (RNeasy Mini Kit Qiagen) with β-mercaptoethanol (10 μl ml^-1^) and 1200 μl 96% (vol./vol.) ethanol were added. For RNA isolation, the RNeasy Mini Kit (Qiagen) was used as recommended by the manufacturer, but instead of RW1 buffer RWT buffer (Qiagen) was used in order to isolate RNAs smaller than 200 nucleotides also. For sequencing, the strand-specific cDNA libraries were constructed with a NEB Next Ultra II Directional RNA library preparation kit for Illumina and the NEB Next Multiplex Oligos for Illumina (New England BioLabs, Frankfurt am Main, Germany). Sequencing was performed by using the HiSeq2500 instrument (Illumina Inc., San Diego, CA, USA) using the HiSeq Rapid SR Cluster Kit v2 for cluster generation and the HiSeq Raid SBS Kit (50 cycles) for sequencing in the single-end mode and running 1×50 cycles. Raw reads have been deposited in the Sequence Read Archive under study SRP451968 (SRR25447477-SRR25447482). More detailed information can be found in the Supplementary Information.

### Conjugation experiments

Conjugation experiments were carried out using strains Teo1 and Teo8 as donors and gentamicin-resistant *P. putida* KT2440::*eyfp-gm* as recipient. Approximately 1 x 10^9^ donor cells and 3 x 10^9^ recipient cells were mixed and spotted onto sterile cellulose acetate filter disks (ca. 3 cm^2^, 0.45 µm pores) on LB agar plates. Conjugation plates were incubated at 30°C for 24 h before cells were washed off the filter disks with 50 mM phosphate buffer (pH 7.2). Decimal dilutions were plated onto minimal medium agar plates with TRIS as only carbon and nitrogen source containing 40 µg/ml gentamicin. Transconjugants were screen using a ChemiDocTM Imaging System (Bio-Rad) using blue epi illumination and an emission filter at 530 nm. The conjugation rate was calculated by dividing the number of positive transconjugants by the number of recipient cells in each assay. The successful transfer of the p1_Teo1 and p2_Teo8 plasmids was confirmed by colony PCR amplifying the *tupA*, *taoA*, *taoB*, *glyA*, and *dsdA* genes (**Suppl. Tab. ST2**).

## Acknowledgment

The authors thank the operators and the employers of the water treatment facilities in the cities of Münster, Hamm, Dülmen, and Güthersloh-Putzhagen for their help with sampling. We especially thank Rudolf Cigelski (Hamm) for his continued support and his great efforts to provide us with samples. We further thank Malte Jochimsen, Anna Hannes, and David Wallbraun for experimental support.

## Competing Interest Statement

The authors declare no conflict of interest.

## Data availability

The whole-genome shotgun projects have been deposited at DDBJ/ENA/GenBank under the BioProject accession numbers PRJNA999090, PRJNA999091, PRJNA999092, PRJNA999094, PRJNA999097, PRJNA999137, PRJNA999141 and the transcriptome data under the BioProject accession number PRJNA999090 with the SRA accession numbers SRR25447477 - SRR25447482.

## Author Contributions

JH conceived the study, designed, planned, and performed experiments, supervised the work, analysed data, and wrote the manuscript. AB performed cloning, conjugation, and isolation experiments and analysed data. LN performed isolation and growth experiment and analysed data. RD contributed reagents and analytic tools. AP performed DNA and RNA extractions, library preparation, DNA sequencing, and transcriptome analysis, and edited the manuscript. BP conceived the study, supervised the work, co-wrote the manuscript, and supplied funding. All authors have approved the final version.

## Competing Interest Statement

The authors declare no conflict of interest.

## Notes

### Competing Interest Statement

The authors have declared no competing interest.

### Summary of Updates

Minor changes in the abstract; changes in the supplemental information; Figure 5 revised

