## Supplemental Information for "A patchwork pathway of apparently recent origin enables degradation of the synthetic buffer compound TRIS in bacteria"

**This PDF file includes:**

Supporting text
Supplemental figures S1 to S5
Supplemental tables ST1 to ST2
Legend for supplemental dataset SD1
References

**Other supporting material for this manuscript include the following:**

Supplemental dataset SD1

**Supporting Information Text**

### Supporting Methods

#### Initial enrichment cultures

A TRIS-buffered artificial wastewater medium (pH 7.0) containing 0.1 mM testosterone, 5 mM TRIS, 3.57 mM $NH_4Cl$, 1.07 mM urea, 0.74 mM $NaNO_3$, 0.12 mM NaCl, 0.036 mM $CaCl_2$ x 2 $H_2O$, 0.017 mM $MgSO_4$ x 7 $H_2O$, 0.0098 mM $K_2HPO_4$ x 3 $H_2O$, 0.0.0042 mM $NaH_2PO_4$ x $H_2O$ and trace elements [1] was used for the initial experiment to enrich testosterone-degrading microorganisms from activated sludge. For solid media plates, 1.5 % agar was added prior to autoclaving. The enrichment culture (35 ml) was inoculated with a fresh active sludge sample (1 ml) from a local wastewater treatment plant (**Supp. Tab. ST1**) and was grown at room temperature in a flow-through chamber containing polyethylene (PE, 4.0 x 4.0 x 0.125 mm) particles to provide a colonization surface for bacteria. When the enrichment cultures showed growth, six overgrown PE particles were transferred to fresh minimal medium and incubated in the same way. This was repeated three times before cultures were spread onto solid minimal medium plates with testosterone and individual colonies were analysed further. The isolate *Pseudomonas hunanensis* strain Teo1 did not degrade the supplied testosterone substrate in liquid cultures but showed consistent growth in the TRIS-buffered artificial wastewater medium.

#### Genome sequencing, assembly, and annotation

Genomic DNA for Illumina sequencing was isolated using the Monarch genomic DNA prep kit (New England Biolabs). High molecular weight DNA (HWD) was isolated with the MasterPure Complete DNA & RNA Purification Kit (Biozym, Hessisch Oldendorf, Germany) as recommended by the manufacturer. Quality of isolated DNA was initially check by agarose gel electrophoresis and validated on an Agilent Bioanalyzer 2100 using an Agilent DNA 12000 Kit as recommended by the manufacturer (Agilent Technologies, Waldbronn, Germany). Concentration and purity of the isolated DNA was first checked with a Nanodrop ND-1000 (PeqLab Erlangen, Germany) and exact concentration was determined using the Qubit dsDNA HS Assay Kit as recommended by the manufacturer (Life Technologies GmbH, Darmstadt, Germany). Illumina shotgun libraries were prepared using the Nextera XT DNA Sample Preparation Kit. To assess quality and size of the libraries, samples were run on an Agilent Bioanalyzer 2100 using an Agilent High Sensitivity DNA Kit as recommended by the manufacturer (Agilent Technologies, Waldbronn, Germany). Concentration of the libraries were determined using the Qubit dsDNA HS Assay Kit as recommended by the manufacturer (Life Technologies GmbH, Darmstadt, Germany). Sequencing was performed on a MiSeq system with the reagent kit v3 with 600 cycles (Illumina, San Diego, CA, USA) as recommended by the manufacturer. For Nanopore sequencing 1.5 µg HWD was used for library preparation using the Ligation Sequencing Kit 1D (SQK-LSK109) and the Native Barcode Expansion Kit (EXP-NBD104 and EXP-NBD114) as recommended by the manufacturer. Sequencing was performed for 72 h using a MinION device Mk1B and a SpotON Flow Cell R9.4.1 as recommended by the manufacturer (Oxford Nanopore Technologies) using MinKNOW software for sequencing and Guppy in high accuracy mode for basecalling and demultiplexing. Unicycler [2] was used with default settings to perform hybrid assemblies. Annotation was performed with Prokka [3] and default settings. Sequencing reads and genome assemblies are available under the project accession numbers PRJNA999090, PRJNA999091, PRJNA999092, PRJNA999094, PRJNA999097, PRJNA999137, PRJNA999141. To resolve the border of the TRIS degradation-encoding region in strain Teo3, DNA was amplified with the primer pairs S/T and S/U (**Suppl. Tab. ST2**) and the PCR products were sequenced using the same primers. Isolated strains were taxonomically classified using the GTDBtk tool (v 1.7.0, [4]) and genome completeness and contamination were assessed using CheckM (v. 1.0.18, [5]) with standard settings.

DNA manipulation and heterologous protein expression

DNA amplification and purification was carried out using standard techniques. DNA for cloning purposes was amplified by PCR using a high-fidelity polymerase (Q5, New England Biolabs). Predicted TRIS degradation genes were cloned into the expression plasmid pUCp18 [6] by isothermal assembly [7]. For this, DNA fragments were PCR-amplified from *P. hunanensis* Teo1 DNA and pUCp18 was linearized using primers with overlapping homologies designed with the NEBuilder online tool (New England Biolabs, Supp. Tab. ST2). Purified fragments and plasmid were combined in equimolar concentrations in isothermal assembly reaction buffer and assembled at 50°C for 60 min. Assembled DNA was transformed into competent *E. coli* cells by chemical transformation and positive clones were selected on LB agar plates containing an appropriate antibiotic. Correct insertion of the fragments into the vector was confirmed by PCR using standard M13 primers and sequencing (Microsynth AG, Switzerland). LB grown overnight *E. coli* cultures carrying predicted TRIS degradation genes on the expression vector pUCp18 were inoculated into fresh LB medium and 0.2 mM (isopropyl β-D-1-thiogalactopyranoside) IPTG were added at an $OD_{600}$ of around 0.4. Cultures were grown to a final $OD_{600}$ of 0.8 – 1.0 before cells were harvested, washed with sterile minimal medium without carbon source, and were resuspended in minimal medium with TRIS to an OD600 of 1.5.

A YFP-tagged *Pseudomonas putida* KT2440 strain was constructed following the protocol from [8] using the plasmid pBK-miniTn7(Gm)PA1/04/03-*eyfp*-a. In short, cells from an overnight culture of P. putida KT2440 were mixed with cells of *E. coli* SM10(λpir) pUX-BF13, *E. coli* JM105 pBK-miniTn7(Gm)PA1/04/03-*eyfp*-a and *E. coli* RK600 and conjugation was carried out at 30°C over night on non-selective LB plates. Cells were washed of the plates with minimal medium without carbon source and were plated onto minimal medium with 10 mM benzoate and gentamycin to select for positive transconjugants. Single colonies showing YFP fluorescence (Chemidoc MP Imaging System, Bio-Rad Laboratories GmbH) were selected and correct insertion of the *yfp*-containing Tn7 element into the chromosome was confirmed by PCR.

If required, kanamycin (20-50 µg/ml), tetracycline (10 µg/ml), or gentamycin (20-40 µg/ml) antibiotics were added to the medium after autoclaving.

<u>Transcriptome analysis</u>

To determine the RNA integrity number the isolated RNA was run on an Agilent Bioanalyzer 2100 using an Agilent RNA 6000 Nano Kit as recommended by the manufacturer (Agilent Technologies, Waldbronn, Germany). Remaining genomic DNA was removed by digesting with TURBO DNase (Invitrogen, Thermo Fischer Scientific, Paisley, UK) The Pan-Prokaryozes riboPOOL kit v4 (siTOOLS BIOTECH, Planegg/Martinsried, Germany) was used to reduce the amount of rRNA-derived sequences. To assess quality and size of the libraries samples were run on an Agilent Bioanalyzer 2100 using an Agilent High Sensitivity DNA Kit as recommended by the manufacturer (Agilent Technologies). Concentration of the libraries was determined using the Qubit dsDNA HS Assay Kit as recommended by the manufacturer (Life Technologies GmbH, Darmstadt, Germany).

For quality filtering and removing of remaining adaptor sequences of the resulting sequences, Trimmomatic-0.39 [9] and a cutoff phred-33 score of 15 were used. The mapping against the reference genomes of Pseudomonas hunanensis Teo1 (BioProject PRJNA999090, Accessions CP131127- CP131129) was performed with Salmon (v 1.9.0) [10]. As mapping back-bone a file that contains all annotated transcripts excluding rRNA genes and the whole genome of the references as decoy was prepared with a k-mer size of 11. Decoy-aware mapping was done in selective-alignment mode with '–mimicBT2', '–disableChainingHeuristic' and '–recoverOrphans' flags as well as sequence and position bias correction and 10,000 bootstraps. For –fldMean and –fldSD, values of 325 and 25 were used, respectively. The quant.sf files produced by Salmon were subsequently loaded into R (v 4.2.0) using the tximport package (v 1.24.0). DeSeq2 (v 1.36.0) [11] was used for normalization of the reads and fold change shrinkages were also calculated with DeSeq2 and the apeglm package (v 1.18.0)

[12]. Genes with a log2-foldchange of +2/-2 and a p-adjust value <0.05 were considered differentially expressed.

Bioinformatic analyses

DNA and protein sequences were queried against the NCBI nucleotide collection (nr/nt) and the non-redundant protein sequence databases, respectively, using the BLASTn and BLASTp algorithms with standard settings. To identify homologs of the TRIS degradation proteins and other proteins encoded on p1_Teo1, the whole p1_Teo1 proteome was used as a query in a best reciprocal BLASTp search [13] against the proteins encoded on plasmids with high similarity to p1_Teo1 or against the whole proteomes of *Pseudomonas sichuanensis* Teo3, *Pseudomonas* sp. strain GXZC, and *Shinella zoogloeoides* Teo12. Similarly, conjugative and replicative proteins were identified on p1_Teo1 by using the respective proteins encoded on pND6-2 as queries in a best reciprocal BLASTp search. Whole plasmid DNA sequences were compared and visualized using progressiveMAUVE alignment (v. 2.4.0). Gene clusters were drawn in R using the gggenes package (v. 0.5.0). Circular plasmids were visualized using Proksee [14]. For this, DNA and proteins were aligned to the p1_Teo1 plasmid using the BLASTn and BLASTp algorithms implemented in Proksee. Orthologs to mobile elements were identified using mobileOG-db (v. 1.1.6) implemented in Proksee. Insertion elements and transposons were identified by BLAST against the ISfinder database using standard settings [15].

For phylogenetic analyses, TaoA and TaoB protein sequences from Teo1 were used as queries for a search against the InterPro database (https://www.ebi.ac.uk/interpro/search/sequence/). For phylogenetic analyses of TaoB all reviewed proteins of the glucose-methanol-choline oxidoreductase family (InterPro IPR012132) were downloaded. For phylogenetic analyses TaoA all reviewed proteins containing an aldehyde dehydrogenase domain (InterPro IPR015590) were downloaded. Proteins were clustered using the easy-cluster algorithm of mmseqs2 [16] with a minimum sequence identity threshold of 0.9 and minimum coverage of 0.8 for TaoB proteins and minimum sequence identity threshold of 0.5 and minimum coverage of 0.8 for TaoA. Protein sequences were aligned using the Muscle algorithm and consensus trees were calculated from maximum likelihood trees with 1000 boostrap repetitions using the W-IQ-TREE online calculator [17]. Trees were visualized with FigTree (v. 1.4.4).

Statistical analysis

Growth, cell suspension, conjugation and transcriptomic experiments were carried out in at least three independent biological replicates. Line graphs were drawn using the mean and error bars denote the standard deviation of the mean.

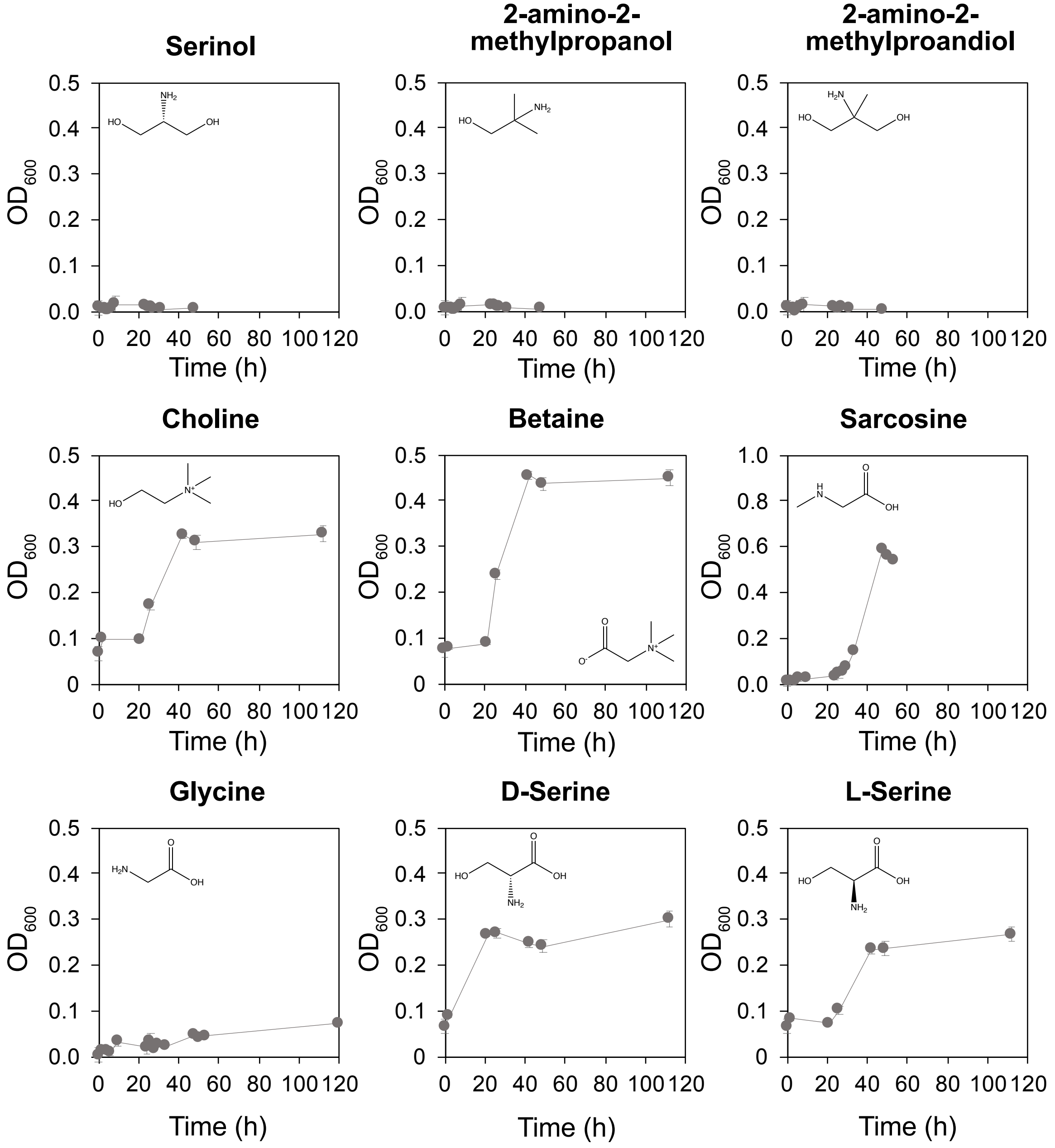


**Fig. S1.** Growth of Teo1 with different substrates. Strain Teo1 was grown with each substrate in three independent replications. Average and standard deviation are shown.

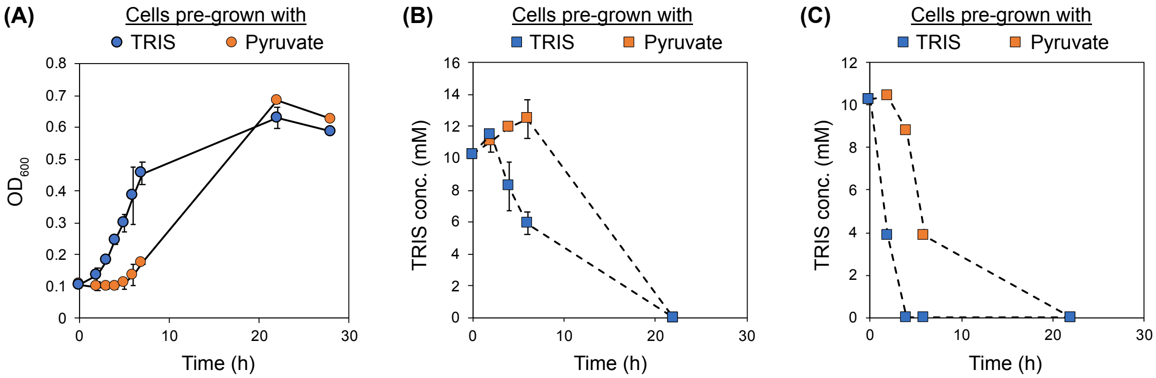


**Fig. S2.** Inducibility of TRIS degradation in strain Teo1 in dependency of prior growth with TRIS. (**A**) Cells that were pre-grown with TRIS as only substrate (blue) immediately started growing when transferred into fresh medium containing TRIS as only substrate. Cells that were pre-grown with pyruvate as only substrate (orange) developed a lag phase of around six hours when transferred to fresh medium containing TRIS as only substrate. (**B**) This is also reflected in a later onset of TRIS degradation in cultures inoculated with cells pre-grown with pyruvate (orange) compared to TRIS pre-grown cells (blue). (**C**) A similar delay in TRIS removal from the medium was observed in high density cell suspensions of strain Teo1 in cells pre-grown with pyruvate (orange) vs. cells pre-grown with TRIS (blue). Each experiment was carried out in three independent replications. Average and standard deviation are shown.

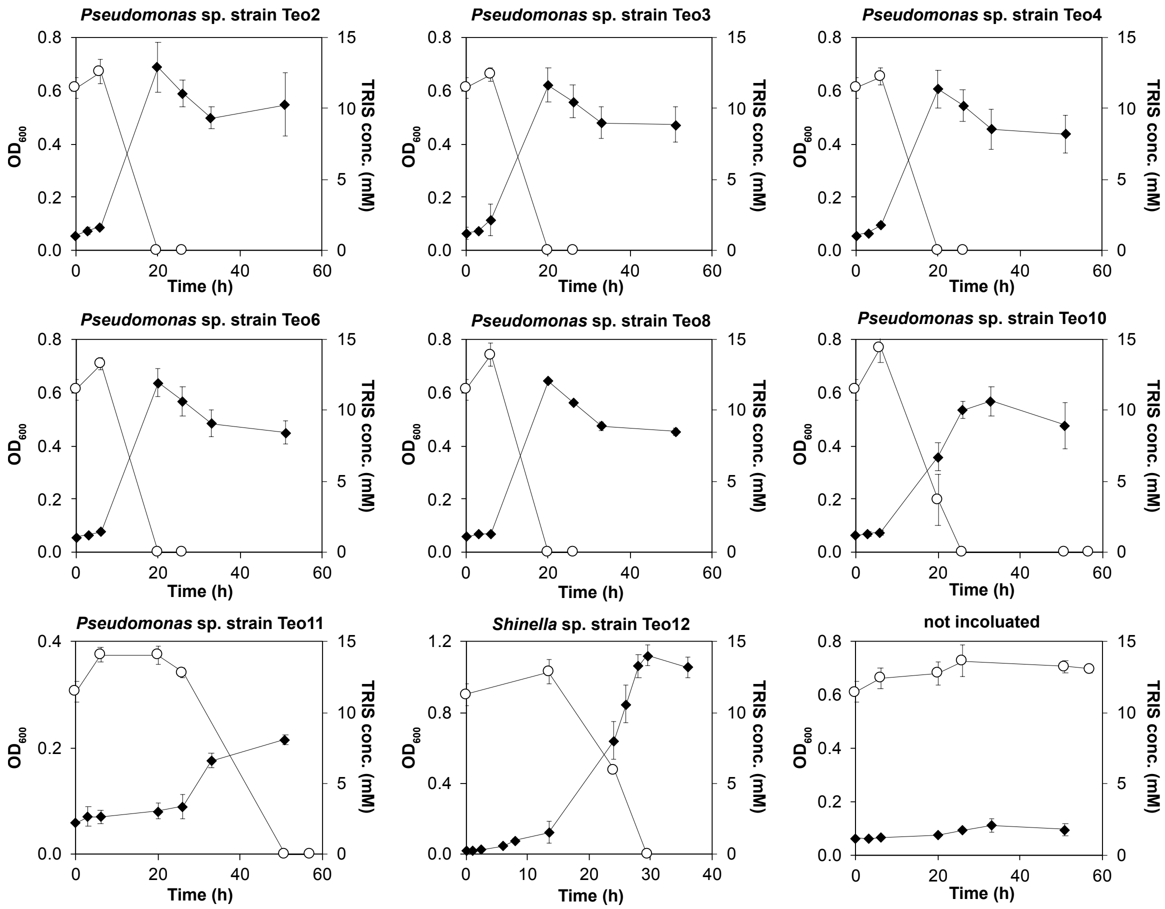


**Fig. S3:** Growth (filled diamonds) and TRIS degradation (open circles) of strains Teo2 – Teo12. Each strain was grown in three independent replications. Average and standard deviation are shown.

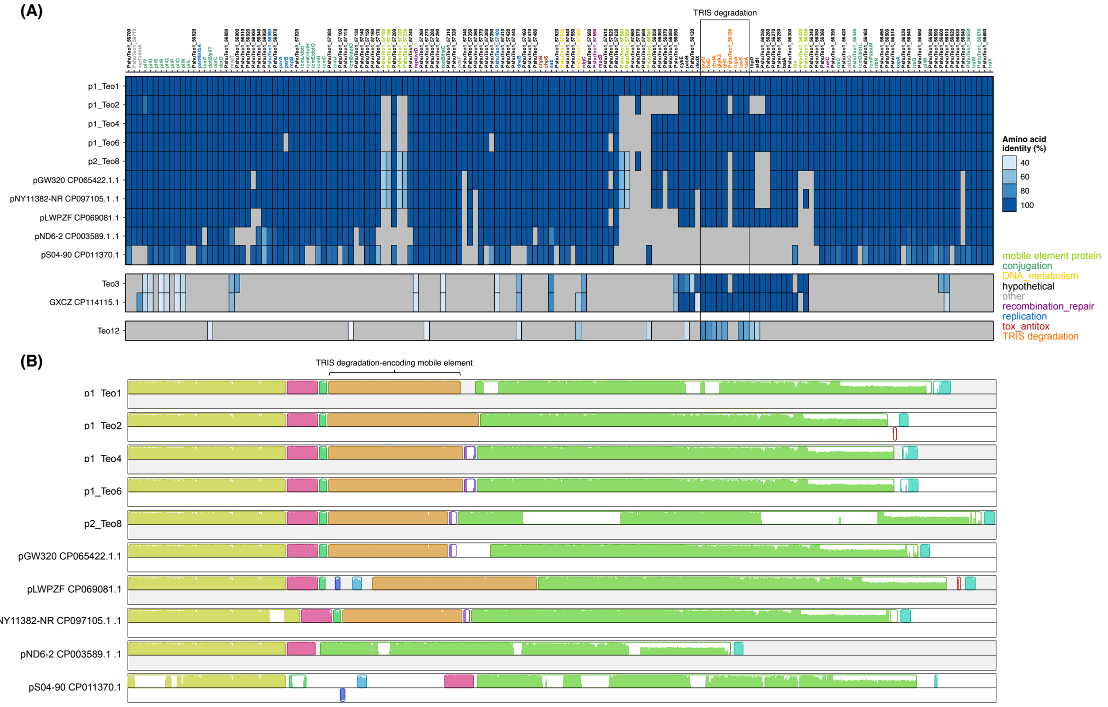


**Fig. S4**: (**A**) Best reciprocal BLASTp analysis with proteins encoded on p1_Teo1 as query against the proteins encoded on similar plasmids identified in this work in TRIS-degrading isolates and plasmids from the NCBI genome database. A separate analysis was also conducted with all proteins encoded in the genomes of Pseudomonas sichuanensis Teo3, Pseudomonas sp. strain GXZC, and Shinella zoogloeoides Teo12. The amino acid identity of homologs of p1_Teo1 proteins encoded on the other plasmids or in the genomes are shown. Grey fields indicate that no homologous protein was found. (**B**) MAUVE alignment of the TRIS degradation-encoding plasmids identified in this study with the closely related plasmids pND6-2 and pS04-90. Coloured blocks indicate regions of the plasmids that are homologous and free from major DNA rearrangement. The height of the similarity profile within each box displays to the average level of conservation in that region of the plasmids.

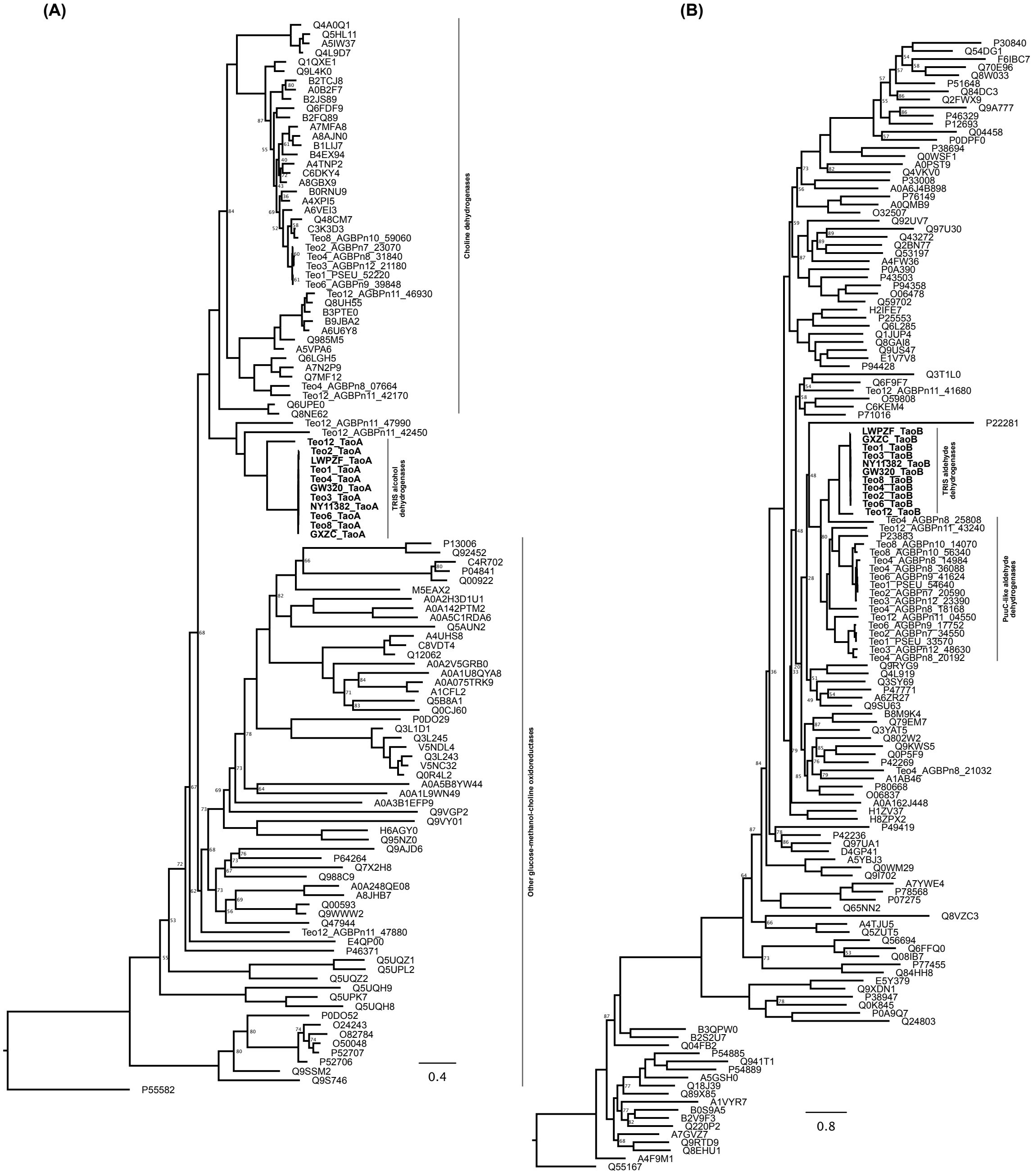


**Fig. S5**: Consensus maximum-likelihood trees of (**A**) TRIS alcohol dehydrogenase proteins (TaoA) with other reviewed proteins from the glucose-methanol-choline oxidoreductase (GMC) family (InterPro IPR012132) and other proteins annotated as choline dehydrogenases in the TRIS-degrading isolates Teo1 – Teo12 and of (**B**) TRIS aldehyde dehydrogenase proteins (TaoB) with other reviewed proteins containing an aldehyde dehydrogenase domain (InterPro IPR015590) and other proteins annotated as PuuC-like aldehyde dehydrogenases in the TRIS-degrading isolates Teo1 – Teo12. Numbers at tree nodes correspond to bootstrap support values (n = 1000) and are not shown for bootstraps values of 90 and higher.

**Table S1.** Isolation sources, taxonomic classification and genome characteristics of TRIS-degrading strain isolated in this study.

| Location (long/lat) | Isolation source | Isolation date | Isolated strain | Taxonomic classification 16S rRNA gene* | Genome completeness and contamination | Taxonomic classification GTDBtk | Genbank Accession |
|---|---|---|---|---|---|---|---|
| Waste water treatment plant (Münster Coerde; 52.00/7.65) | Activated sludge | Jan 17, 2020 | **Teo1** | *Pseudomonas putida* | compl. 100.00 %, cont. 0.29 % | *Pseudomonas hunanensis* | |
| Waste water treatment plant (Münster Coerde; 52.00/7.65) | Actvated sludgde | Oct 03, 2020 | **Teo2** | *Pseudomonas putida* | compl. 94.28 %, cont. 0.43 % | *Pseudomonas hunanensis* | |
| Water purification plant (Münster Hornheide; 51.88/7.68) | Activated charcoal | Oct 03, 2020 | **Teo3** | *Pseudomonas sichuanensis* | compl. 97.56 %, cont. 1.23 % | *Pseudomonas sichuanensis* | |
| Lake Aasee (Münster; 51.94/7.59) | Water sample | Oct 03, 2020 | | **no isolate** | | | |
| City pond (Münster; 51.97/7.62) | Water sample | Oct 03, 2020 | | **no isolate** | | | |
| City pond (Münster; 51.97/7.60) | Water sample | Oct 03, 2020 | | **no isolate** | | | |
| Waste water treatment plant (Hamm West; 51.67/7.73) | Activated sludge | Jan 15, 2021 | **Teo4** | *Pseudomonas* sp. | compl. 97.39 %, cont. 1.56 % | *Pseudomonas* sp. | |
| Waste water treatment plant (Hamm Mattenbecke; 51.69/7.82) | Activated sludge | Jan 15, 2021 | **Teo6** | *Pseudomonas asiatica* | compl. 100.00 %, cont. 0.35 % | *Pseudomonas asiatica* | |
| Waste water treatment plant (Hamm Uentrop; 51.69/7.94) | Activated sludge | Jan 15, 2021 | **Teo8** | *Pseudomonas extremaustralis* | compl. 100.00 %, cont 1.35 % | *Pseudomonas extremaustralis* | |
| Lake Aasee (Münster; 51.94/7.59) | Water sample | Jun 18, 2022 | **Teo10** | *Pseudomonas* sp. | not sequenced | | |
| Waste water treatment plant (Dülmen; 51.81/7.26) | Activated charcoal | Jun 18, 2022 | **Teo11** | *Pseudomonas extremaustralis* | not sequenced | | |
| Waste water treatment plant (Dülmen; 51.81/7.26) | Activated charcoal | Jun 18, 2022 | **Teo12** | *Shinella zoogloeoides* | compl. 99.48 %, cont. 0.19% | *Shinella zoogloeoides* | |
| Waste water treatment plant (Dülmen; 51.81/7.26) | Activated sludge | Jun 18, 2022 | | **no isolate** | | | |
| Waste water treatment plant (Putzhagen; 51.90/8.34) | Activated charcoal | Jun 18, 2022 | | **no isolate** | | | |
| Rieselfelder (Münster; 52.03/7.65) | Water sample | Jun 18, 2022 | | **no isolate** | | | |
| City pond (Münster; 51.96/7.61) | Water sample | Jun 18, 2022 | | **no isolate** | | | |

* taxomonic classification based on majority of taxonomic assignments of the 10 best hits against the NCBI nr nucleotide database **completeness and contamination were determined with CheckM (v1.0.18)

**Table S2.** Primers used in this study.

| # | Primer | Sequence | Function |
|---|---|---|---|
| A | tupA_gibson_fw | CACAGGAAACAGCTATGACCAACTACAGGAAGCTTAAACGATGAATC | fw primer gibson assembly |
| B | taoB_gibson_fw | CACAGGAAACAGCTATGACCTCCAGCAATGAGGAAACAGCATGACC | fw primer gibson assembly |
| C | tupA_gibson_rev | GTTGTAAAACGACGGCCAGTTTACTCGCAGAGCACCGAC | rev primer gibson assembly |
| D | taoA_gibson_rev | GTTGTAAAACGACGGCCAGTTCAGTGGGGCATGCCGGC | rev primer gibson assembly |
| E | pUCp18_gibson_fw | ACTGGCCGTCGTTTTACAACGT | fw primer gibson assembly |
| F | pUCp18_gibson_rev | GGTCATAGCTGTTTCCTGTGTG | rev primer gibson assembly |
| G | tupA_fw | CAATGCCTAGAGCAACACGC | fw primer to amplify tupA gene |
| H | tupA_rev | GAAGGTCTTTCCGTCCAGGG | rev primer to amplify tupA gene |
| I | taoB_fw | GCGCTTGTGTTCATCGTGTT | fw primer to amplify taoB gene |
| J | taoB_rev | ACGAAGACCATGCCGTTGAT | rev primer to amplify taoB gene |
| K | taoA_fw | GGCTACTACATCGAGCCGAC | fw primer to amplify taoA gene |
| L | taoA_rev | CGAAAGCCAGCTGGGAAAAG | rev primer to amplify taoA gene |
| O | glyA_fw | TGGCTTTCAGGTTCGGGTTT | fw primer to amplify glyA gene |
| P | glyA_rev | TCTCTCAAGGCACGCAACTT | rev primer to amplify glyA gene |
| Q | dsdA_fw | ACGGTCGGCCAAGAATTGAT | fw primer to amplify dsdA gene |
| R | dsdA_rev | AAACCCGATTCCGTTCGACA | rev primer to amplify dsdA gene |
| S | Teo3_cont3_fw | TCGAGTCAGTGTTTCCCGTG | fw primer binding to Teo3 contig 3 |
| T | Teo3_cont5_rev_a | AGACGTAATTGCCGGTCAGG | rev primer binding to Teo3 contig 5 |
| U | Teo3_cont5_rev_b | GGGCGAATAGAGAAAACGGA | rev primer binding to Teo3 contig 5 |

**Dataset SD1 (separate file).** Transcript abundances of genes of *Pseudomonas hunanensis* Teo1 grown with TRIS (PshuTeo11 - PshuTeo13) or pyruvate (PshuTeo14 - PshuTeo16). DeSeq2 was used to calculate $log_2$ fold changes and p values.
